# Behavioral flexibility is associated with changes in structure and function distributed across a frontal cortical network in macaques

**DOI:** 10.1101/603530

**Authors:** Jérôme Sallet, MaryAnn P Noonan, Adam Thomas, Jill X O’Reilly, Jesper Anderson, Georgios K Papageorgiou, Franz X Neubert, Bashir Ahmed, Jackson Smith, Andrew H Bell, Mark J Buckley, Léa Roumazeilles, Steven Cuell, Mark E Walton, Kristine Krug, Rogier B Mars, Matthew FS Rushworth

## Abstract

One of the most influential accounts of central orbitofrontal cortex– that it mediates behavioral flexibility – has been challenged by the finding that discrimination reversal in macaques –the classic test of behavioral flexibility –is unaffected when lesions are made by excitotoxin injection rather than aspiration. This suggests the critical brain circuit mediating behavioral flexibility in reversal tasks lies beyond the central orbitofrontal cortex. To determine its identity a group of nine macaques were taught discrimination reversal learning tasks and its impact on grey matter was measured. Magnetic resonance imaging scans were taken before and after learning and compared with scans from two control groups each comprising ten animals. One control group learned similar discrimination tasks but which lacked any reversal component and the other control group engaged in no learning. Grey matter changes were prominent in posterior orbitofrontal cortex/anterior insula but also were found in three other frontal cortical regions: lateral orbitofrontal cortex (12o), cingulate cortex, and lateral prefrontal cortex. In a second analysis, neural activity in posterior orbitofrontal cortex/anterior insula was measured at rest and its pattern of coupling with the other frontal cortical regions was assessed. Activity coupling increased significantly in the reversal learning group in comparison to controls. In a final set of experiments we used similar structural imaging procedures and analyses to demonstrate that aspiration lesion of central orbitofrontal cortex, of the type known to affect discrimination learning, affected structure and activity in the same frontal cortical circuit. The results identify a distributed frontal cortical circuit associated with behavioral flexibility.

## Introduction

One of the most influential accounts of orbitofrontal (OFC) function suggests it mediates behavioral flexibility in response to changes in the environment [1–4]. Typically, this has been assessed with discrimination reversal (DisRev) learning tasks in which animals learn one choice leads to reward while another does not. Usually the correct and incorrect choices are defined as the selection of one stimulus rather than another but sometimes spatially defined choices are employed. Animals learn to make the reward-associated choice but once they make it reliably the reward assignments are switched so that the previously unrewarded choice becomes the only one followed by reward. Central OFC lesions centered on the orbital gyrus (areas 13 and 11) have long been thought to impair DisRev and the activity of OFC neurons has been related to key DisRev events [1, 4–8].

The consensus view of OFC function has recently been questioned by the finding in macaques that while DisRev is impaired by aspiration lesions of central OFC, it is unimpaired if the lesions are made with excitotoxic injections sparing fibers of passage [3]. The implication is that behavioral flexibility in DisRev performance depends on a specific network of brain regions that are disconnected from one another by aspiration lesions of OFC (which may damage fibers of passage) but not by excitotoxic lesions. The identities of the components of this circuit are, however, unknown.

One key region of interest is the cortex lying laterally adjacent to the central OFC. This has sometimes been referred to as lateral OFC (lOFC) and corresponds to the orbital part of area 12, 12o. This region is important for the linking of a choice to an outcome and using knowledge of such linkages to guide behavior. For example activity here reflects the use of a win-stay/lose-shift behavioral strategy [9]. Win-stay/lose-shift strategies require specific outcome events to be linked to specific choices which are then repeated or avoided in the future depending on how successful they have been. The corresponding region in the human brain [10] has been linked to the learning of specific choice-reward associations [11–13]. Decisions are no longer driven by knowledge of the causal relationships between choices and outcomes when lesions are made that include this region and adjacent ventrolateral prefrontal cortex in the Rhesus macaque [14, 15].

Another potentially important candidate region is the posterior OFC and adjacent anterior insula (AI); Rudebeck and colleagues [3] showed that small aspiration lesions nearby, placed across the posterior OFC, were sufficient to impair DisRev. However, one interpretation of the finding is that damage to white matter pathways in the vicinity, such as the uncinate fascicle, extreme capsule, and cingulum bundle [16, 17] cause the DisRev impairment. Many prefrontal regions, including posterior OFC and AI, are interconnected by these pathways.

An additional reason for thinking that AI and posterior OFC might be important for DisRev is that OFC lesions in new world monkeys and rats, even when made by excitotoxin injection, have been reported to cause DisRev impairments [4, 8, 18–21]. It is possible that the regions referred to as OFC in rodents and new world monkeys have similarities with the posterior OFC and adjacent AI of old world primates such as macaques [22, 23]. Secondly, Wittmann and colleagues (in preparation) have recently shown that activity in this region of the macaque brain reflects not just the outcome of the last choice made but also a longer term history of reward. An animal learns to balance the weight of influence exerted by these signals when it becomes proficient at performing DisRev.

Additional candidate regions within the network of areas thought to be important for reward-guided decision making and which have connections that might be compromised by the white matter damage likely to be associated with an OFC aspiration lesion include the anterior part of the cingulate cortex involved in tracking key features of the reward environment [24–28] and holding the value of alternative or counterfactual choices in order to guide changes in behavior [29–31].

To identify the wider network of brain regions involved in DisRev learning we carried out a series of behavioral, lesion, and neuroimaging experiments in macaques and examined changes in grey matter and functional connectivity. It is known that learning of related tasks in rodents is accompanied by plastic changes in long range axons in frontal cortex [32] and it is known that structural and functional imaging techniques can identify candidate regions in which such changes may occur [33].

First, in experiment 1, we carried out a longitudinal experiment in which we sought brain regions where structural changes were associated with DisRev learning in a group of nine macaques. Structural magnetic resonance imaging (MRI) scans under anaesthesia were taken before and after DisRev learning and compared with matched scans from two groups of control animals either taught to perform a similar reward-guided discrimination task that lacked any reversal component (ten animals) or no task at all (ten animals). Next, to identify regions affected by transneuronal degeneration caused by central OFC aspiration lesions, we used similar techniques to determine whether changes occurred in the same neural circuit when non-fiber sparing lesions were made in central and medial OFC in two macaques. Grey matter throughout the brain in these two animals was compared with grey matter throughout the brain in MRI scans from 28 control macaques. Finally, in experiments 3 and 4, we examined changes in functional activity within the areas identified in experiments 1 and 2. Equivalent analyses were performed in functional MRI (fMRI) scans taken before and after learning (experiment 3) as well as in lesion and control animals (experiment 4). We sought converging evidence for brain regions both affected by the aspiration lesion of OFC in experiment 2 and in which structure and activity were modulated by DisRev learning in experiments 1 and 3, reasoning that effects that replicated across the different studies would be the most reliable.

## Results

### Experiment 1

First, we investigated the wider structural network of brain regions affected by DisRev learning. We sought brain regions where structural (experiment 1) and activity (experiment 3) changes were associated with DisRev experience in a group of nine macaques. Four animals chose between stimuli with different identities (Object DisRev) and five chose between different target locations (Spatial DisRev). We obtained structural MRI and functional MRI (fMRI) measures at two time points before and after behavioral training (pre and post learning) while animals were under general anesthesia (fig.1). At Scan 1 the animals had some experience with the experimental apparatus and had learned that pressing a target on either the left or right of a touchscreen was associated with a juice reward. By contrast, at Scan 2 animals had substantial experience of reversal tasks and could perform five reversals per day, every time 25 correct choices had been made, at high levels of accuracy (≥80% correct on average throughout the entire session). Learning has previously been associated with structural and activity changes that can be measured with MRI even when subjects are at rest [33]. Typically, such investigations have focused on sensorimotor learning and consequently on sensorimotor brain regions but given the connectional changes observed in rodents during learning [32] there is no reason why the learning of a “cognitive set” for behavioral flexibility may not be associated with similar structural and activity changes in other brain regions.

**Figure 1.**
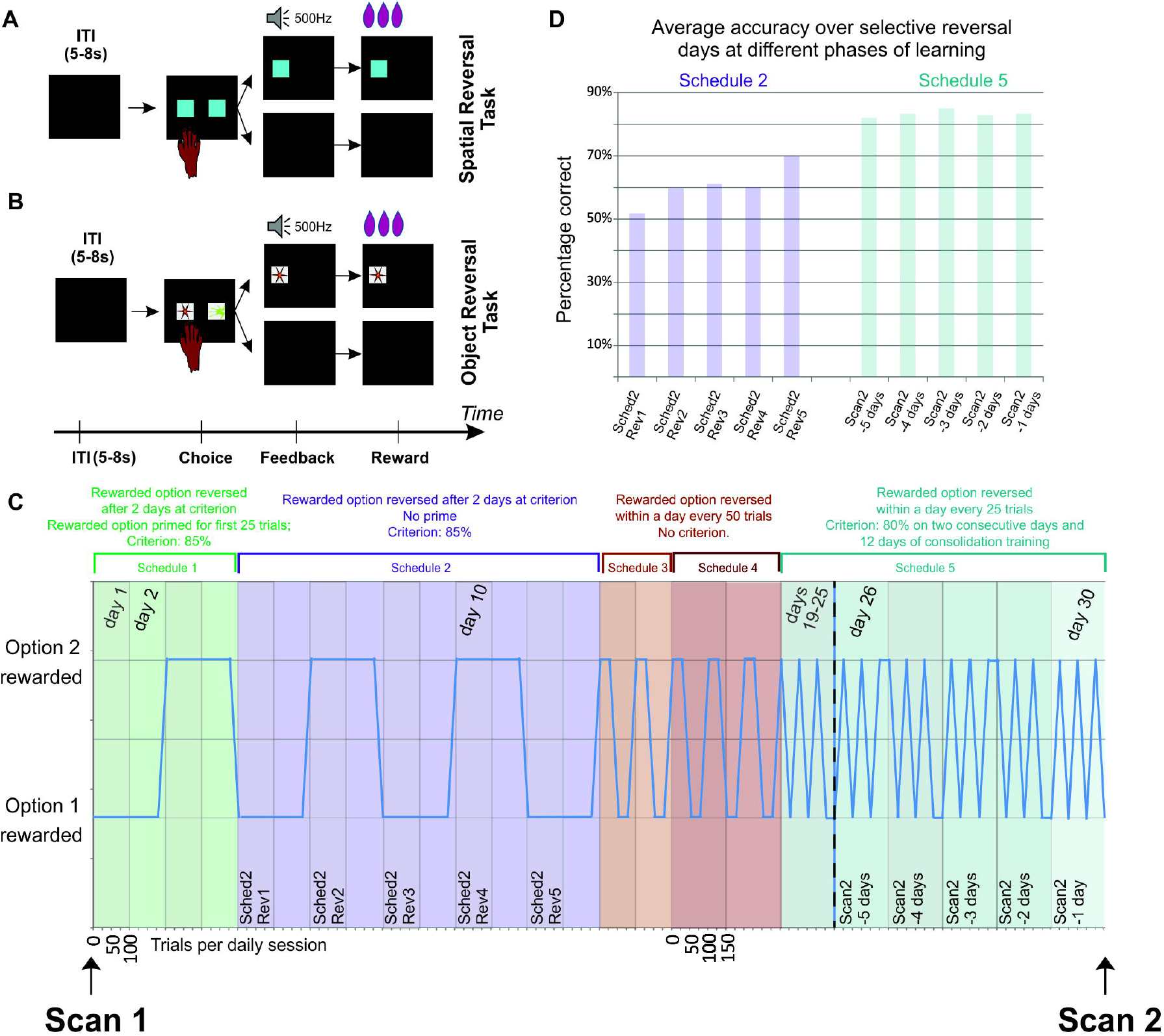
Trial structure in **(A)** Spatial DisRev **(B)** Object DisRev and **(C)** overview of DisRev experience between structural scan 1 and 2 in a longitudinal investigation of the effect of learning on brain structure and function. The schematic illustrates an ideal performer. The minimum number of trials needed by an ideal performer per daily session are illustrated on the abscissa and reward properties of the two options are plotted on the ordinate. In the first testing schedules, animals completed 100 trials per days and one of two simultaneously presented options was designated the rewarded target for the day. In schedule 1 the rewarded choice was primed for 25 trials before introducing, in addition, the unrewarded choice on the opposite side of the screen for an additional 75 trials. The choice-reward contingencies were reversed the day after each animal performed above 85% correct for two consecutive days. Schedule 2 was similar but now the rewarded choice was not primed. Animals experienced five reversals under schedule 2. With perfect performance an animal could, therefore, complete this second phase in ten days. Schedules 3 and 4 introduced a choice-reward association reversal within a given day’s testing session; the reversal occurred once the animal had correctly chosen the target 50 times. The animals completed two days of 100 trials and three days of 150 trials on schedule 3 and 4 respectively. There was no performance criterion in these phases. In the final schedule, the rewarded targets reversed after 25 correctly performed trials. In total the animals had to perform 150 correct trials in a day. Their second scan took place once performance exceeded 80% correct for two consecutive days, and after 12 subsequent days of consolidation training. **(D)** Average correct performance over session for the 1^st^ 5 sessions they encountered a reversal (Schedule 2 sessions; see Fig. 1C) and for the last 5 days prior to the second scan (Schedule 5 sessions). During Schedule 2 monkeys performed at approximately 55% accuracy but by Schedule 5 reversal they performed at 80% accuracy.

A GLM was used to compare grey matter in the nine DisRev learning macaques at scan 1 and scan 2 (mean 0.71 yrs, 0.28std) with that in similarly spaced scans (mean 0.98 years, 0.76std) in 20 controls macaques that lacked experience of DisRev (Supplementary Table 1). Half the control animals (n=10) had no experience of formal training (NoDis Control), while the other half (n=10) had experience of discriminating visual stimuli but in the absence of any reversal requirement (Dis Control). There was no significant difference in the number of days between scan 1 and scan 2 when DisRev learners and All Controls were compared (t_27_ = 1.03, p = 0.312). Structural MRI data were submitted to a deformation based morphometric (DBM) analysis using the Oxford Centre for Functional Magnetic Resonance Imaging (FMRIB) Software Library (FSL) tools FNIRT and Randomise [34] (Methods and Supplementary Methods for details). The logic of the approach is that if a group of brain images can be warped to an identical image, then volumetric changes involved in that warping process give measures of the local differences in brain structure between individuals. Related analyses have previously been described [35, 36]. In addition to our regressor of interest (Behavioral condition: DisRev learners, NoDis Control, Dis Control) we also included control regressors indexing the age and sex of individual monkeys.

At each voxel in the brain the dependent variable was the determinant of the Jacobian matrix from the non-linear registration of each individual’s structural MRI to the group average brain. This is a scalar variable representing how much each voxel in an individual’s brain would need to be expanded or compressed to match the group average brain. To check the reliability of our findings we sought regions in which effects were replicated in both hemispheres by testing for the conjoint probability of symmetrical effects in the two hemispheres with a p<0.001 (under the null hypothesis effects are expected to be randomly distributed across hemispheres [37]) and constituted by more than 15 contiguous voxels (Supplementary Methods for details). In examining the bilaterality of our effects, we adopt an approach suggested for MRI voxel-based grey matter analyses. It is based on the principle that taking into account the spatial extent, across adjacent MRI voxels, of any statistical effect is not necessarily appropriate for grey matter analyses [38] and so alternative tests of robustness have been advocated that involve examining whether effects are bilaterally symmetrical [35, 39, 40]. The premise rests on the assumption that if a statistical effect noted had a chance of occurrence of p=0.05 in one brain area under the null hypothesis, then it has the chance of occurring in the same area in both hemispheres with the square of this probability (e. g. p=0.05×0.05=0.0025). As explained above, here we report bilaterally symmetrical effects with a conjoint probability of a p<0.001.

Significant changes in grey matter in several regions were observed (fig. 2A, Supplementary Table 2). We focus here on those regions of significant change that were also identified as important in the very different approach undertaken in experiment 2. Significant grey matter increases were associated with DisRev experience compared to All Controls in four parts of the frontal lobes including the anterior part of a region we have previously referred to as lateral orbitofrontal cortex (lOFC) [9–11] but which is more unambiguously identified by the term 12o, posterior lateral OFC extending into anterior insular (plOFC/AI), ventral bank of the principal sulcus in the lateral prefrontal cortex (lPFC), and in the anterior part of cingulate cortex (ACC) close to the region referred to as midcingulate cortex (MCC) by Procyk and colleagues [41]. We refer to it here as ACC/MCC. For illustration we present the averaged residual Jacobian values extracted from bilateral ROIs placed at the center of gravity of the significant clusters.

**Figure 2.**
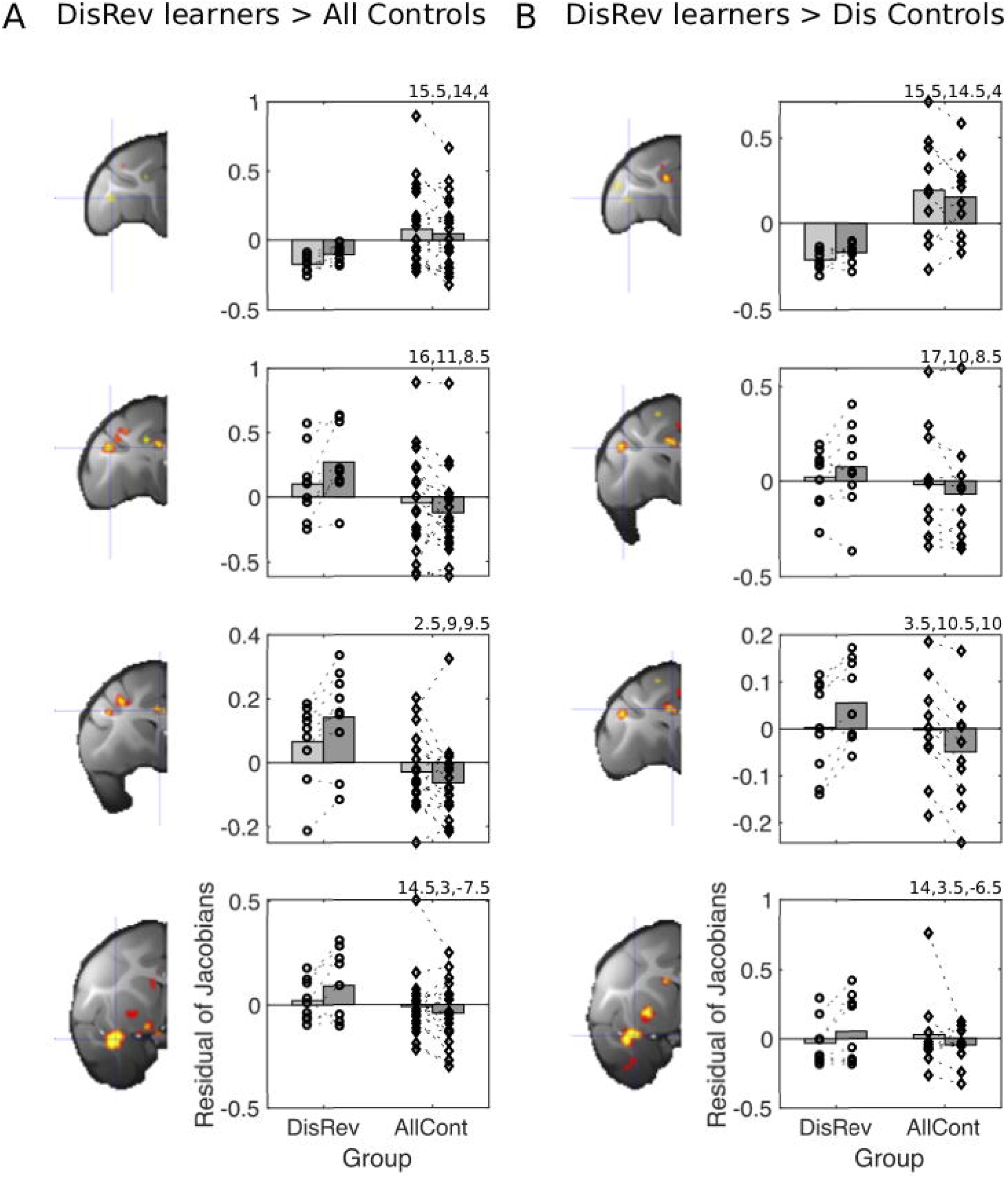
Learning DisRev was associated with distributed grey matter increments **(A)** relative to all control animals including in the same set of brain regions as identified experiment 2: lOFC/12o (cross hairs in top row), ventral bank of the principle sulcus in anterior lPFC (cross hairs in second row), ACC/MCC (cross hairs in third row), and plOFC/AI (cross hairs in bottom row). **(B)** Illustrates that excluding NoDis Controls from the analysis revealed the same set of brain regions with increases in grey matter in lOFC/12o, ventral bank of the principle sulcus, ACC/MCC, and plOFC/AI. Results of analysis testing for bilaterally symmetrical effects (p < 0.001) are shown in yellow and red (p < 0.005). Because in the initial stages of analysis, all scans from both before and after training are registered to a template derived from their group average, the baseline residual Jacobian values in each figure lie close to the mean.

Further post-hoc tests examined whether effects reflected increases in grey matter in DisRev learners or decreases in controls. A Session (two levels: Scan 1-versus Scan 2) x Area (four levels: lOFC/12o, plOFC/AI, lPFC, ACC/MCC) x Hemisphere (two levels: right and left) x Structural scan (two levels) analysis found grey matter across ROIs tended to increase in learners (main effect of Session: F_1,8_= 19.51, p=0.002) while reducing in all controls (main effect of Session: F_1,19_=5.26, p=0.033).

To confirm that the effects were not dependent on non-specific effects associated simply with learning to discriminate between visual stimuli, as opposed to the specific effect of reversing reward contingencies, we examined the contrast between the DisRev learners and Dis Control (and excluded the NoDis Control animals). Significant grey matter increases were notably associated with DisRev experience bilaterally (p<0.001) in the same regions (lOFC/12o, plOFC/AI, lPFC, ACC/MCC, (fig.2B; see Supplementary table 3). Again, we illustrate the averaged residual Jacobian values extracted from bilateral ROIs placed at the centers of gravity of the significant clusters. In summary, changes in these four regions – lOFC/12o, plOFC/AI, lPFC, and ACC/MCC – reflect processes common to DisRev learning regardless of whether choices are defined by stimulus features or spatial position.

### Experiment 2

In order to investigate this network further and identify the wider structural network associated with central OFC aspiration lesions, we looked at the impact on grey matter across the brain of OFC lesions extending from the rostral sulcus on the medial surface of the frontal lobe to the medial bank of the lateral orbital sulcus in 2 macaques. The lesions included tissue sometimes described as ventromedial prefrontal cortex (vmPFC) or medial orbitofrontal cortex (mOFC) [24, 42] as well as the central parts of OFC (fig. 3A). However, they spared tissue lateral to the lateral orbitofrontal sulcus implicated in linking choices and outcomes and the use of win-stay/lose-switch strategies during learning [9, 14]. Thus, the lesion was focused on areas 14, 11, and 13 but spared area 12 including the orbital part of 12, 12o [43, 44]. We refer to the lesions as vmPFC/OFC lesions. We collected structural MRI scans from two macaques approximately 4 months after the aspiration lesions (fig. 3A) and compared them with structural MRIs from a control group of 28 control macaques (Supplementary Table 1). Note, that the control group comprised some of the animals that would go on to learn the DisRev task in experiment 1 but all the scans examined were the ones taken at the first time point prior to any DisRev learning. The large control group made the DBM analysis sensitive to subtle changes in grey matter distant from the primary intended lesion site without requiring a large lesion group. The post-lesion delay in scanning was used to ensure that any neural changes were specific to the lesion intervention and not reflective of any general, immediate consequence of edema beyond the immediate lesion site that might occur during the immediate post-surgery recovery period. In addition to our regressor of interest (control versus lesion group), the age and sex of individual monkeys were included as control regressors in the general linear model (GLM). As in experiment 1, the dependent variable was the determinant of the Jacobian matrix from the non-linear registration of each individual’s structural MRI to the group average brain and again we sought regions in which effects were replicated in both hemispheres by testing for the conjoint probability of symmetrical effects in the two hemispheres with a p<0.001 extended over more than 15 contiguous voxels (Supplementary material). We focused on regions of grey matter reduction in the lesion group relative to controls.

**Figure 3.**
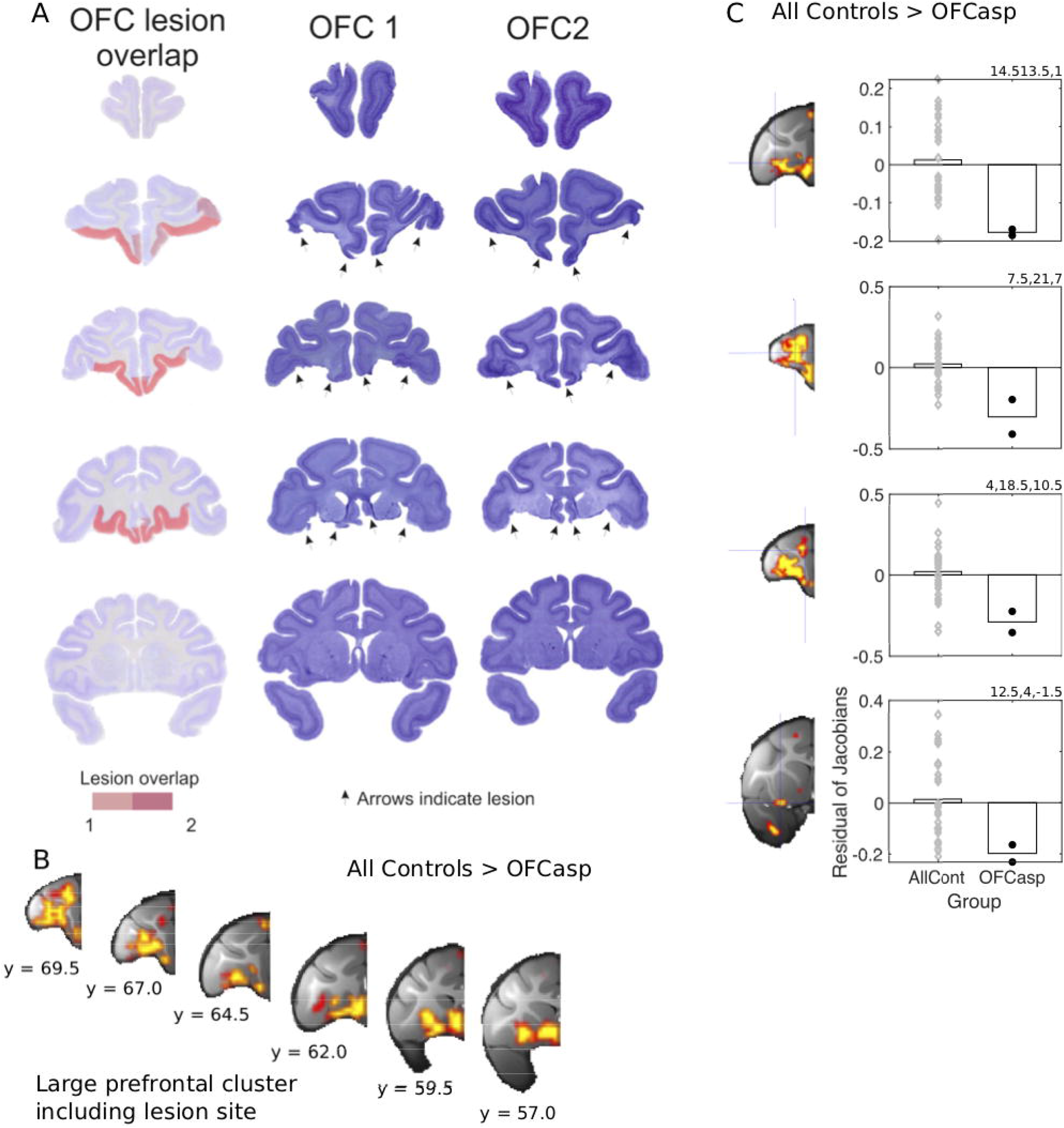
Lesions of central and medial OFC (OFC) and adjacent ventromedial prefrontal cortex (vmPFC) between the lateral orbital sulcus and the rostral sulcus. **(A)** shows coronal sections through the frontal cortex (the most rostral section is shown at the top left and the most caudal on the bottom right). The lesions were intended to cover the same region investigated by Rudebeck and colleagues [3]. The actual lesion is indicated by red coloring. Saturated red color indicates area of lesion overlap in both animals while paler red color indicates area of lesion in a single individual. Lesions were associated with large grey matter decrements in a distributed set of brain regions including not only the aspiration lesion site itself **(B)**, but also the four frontal regions associated with DisRev learning in experiment 1 including lOFC/12o (top row), lPFC on the ventral bank (second row), ACC/MCC, and plOFC/AI **(C)**. Results of analysis testing for bilaterally symmetrical effects (p < 0.001) are shown in yellow and red (p < 0.005). Because in the initial stages of analysis, all scans from both before and after training are registered to a template derived from their average, the baseline residual Jacobian values in each figure lie close to the mean.

Aspiration lesions of vmPFC/OFC were associated with significant changes in grey matter in the frontal cortical regions identified in experiment 1: lOFC/12o, plOFC/AI, ventral bank of the principal sulcus (lPFC) as well as the ACC/MCC (fig. 3B,C). Changes were also noticeable in some other areas (Supplementary Table 4).

For illustrative purposes we present the average Jacobian values, after age and sex had been accounted for, in the lesion group and controls. Jacobian values were extracted from bilateral ROIs (3.375 mm^3^ in diameter) at coordinates reflecting the center of gravity of the bilateral cluster. In some cases, the effects were in relatively adjacent and large frontal clusters (lOFC/12o; lPFC). Therefore, to ensure that Jacobian values extracted from each ROI were independent of one another they were taken from ROIs centered on coordinates just off the center of gravity of lOFC/12o and lPFC. Thus, each illustration reflects separate data as much as possible while still representing the anatomical regions. The precise coordinates of the ROI centers are all reported (Supplementary Table 4).

### Experiment 3

To further characterize the regions identified in Experiments 1 and 2, we examined functional connectivity of the plOFC/AI where grey matter effects associated with DisRev training were greatest and most extensive in experiment 1 and where grey matter effects overlapped in experiments 1 and 2. The blood oxygen level dependent (BOLD) signal was measured under anaesthesia with fMRI. We examined whether pattern of coupling of the activity in plOFC/AI changed after DisRev learning (experiment 3) and after vmPFC/OFC lesions (Experiment 4).

We examined the whole brain functional connectivity of ROIs (15.625mm^3^) in bilateral plOFC/AI (Fig. 4A) using a seed-based correlation analyses in which the GLM design was equivalent to that described in Experiment 1. However, rather than examining grey matter changes associated with DisRev learning, now the focus was on changes in the coupling of fMRI-measured activity. A within-subject, repeated measures design compared the functional connectivity of the bilateral plOFC/AI ROI (fig. 4A) at the time of Scan 1, prior to learning, and Scan 2, after learning in DisRev animals and all Control animals (n=14). Resulting voxelwise p-maps were small volume cluster-corrected using threshold-free cluster enhancement methods [45] (p < 0.05) using large anatomical masks focused on the frontal regions identified across Experiment 1 and 2; namely lOFC/12o, lPFC and ACC/MCC.

**Figure 4.**
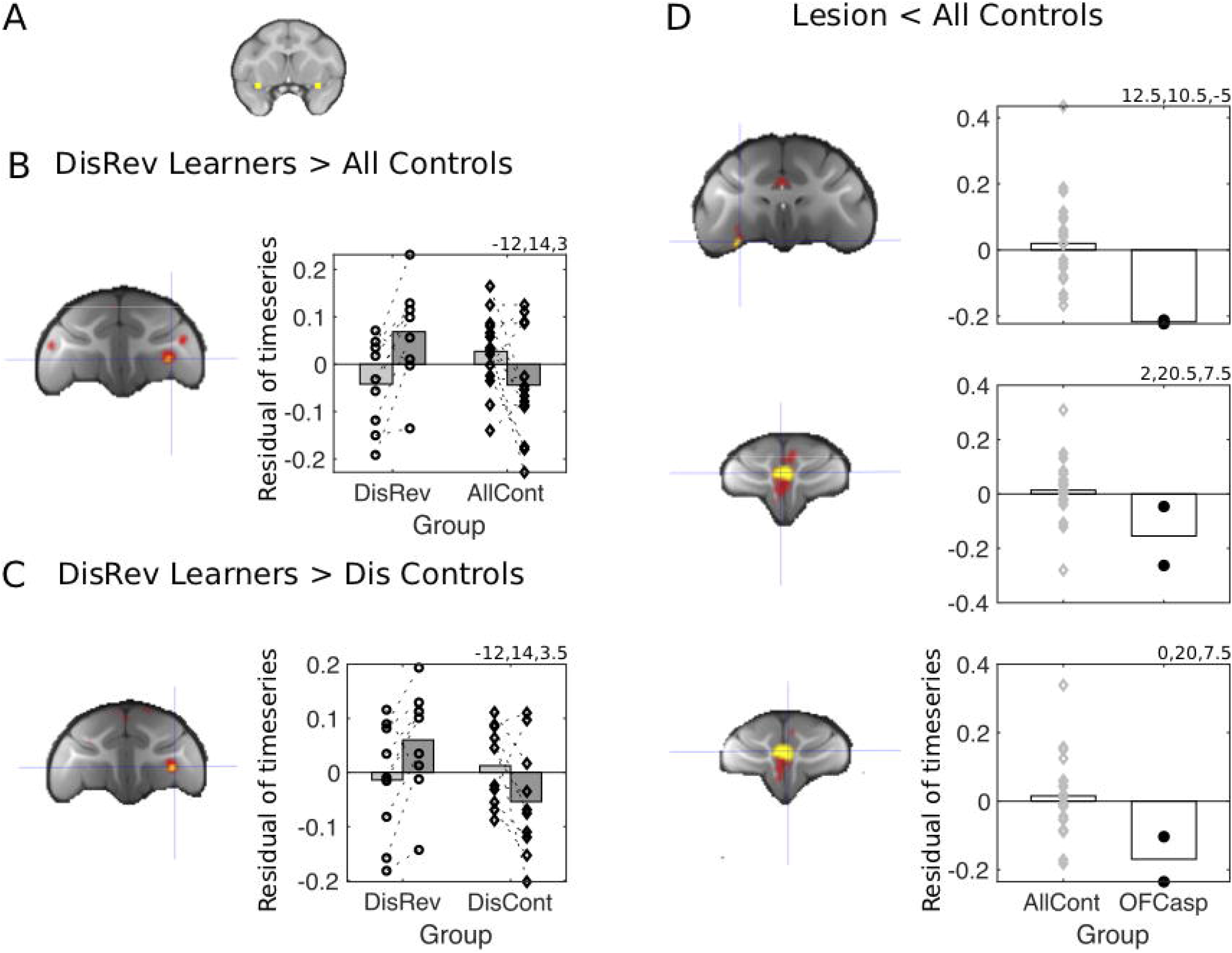
fMRI-measured activity coupling, at rest, between the brain regions identified in Figure 2 and 3. Learning DisRev, relative to all control animals, was associated with changes in functional coupling seeded bilaterally within the plOFC/AI **(A)** within some of the same set of brain regions as experiment 1: **(B)** plOFC/AI increased in functional coupling with the lOFC/12o. Results of Threshold-Free Cluster Enhancement (tfce) correction (p < 0.05) are shown in yellow and red (p < 0.01). **(C)** When the analysis was repeated but now the comparison was between DisRev learners and the Dis Control group that had learned a discrimination learning task lacking a reversal component, the same results was found. **(D)** fMRI functional coupling of the plOFC/AI in the OFC aspiration lesion animals versus controls. Functional coupling between the plOFC/AI and lOFC/12o (top) and clusters in the left and right ACC/MCC identified in the ROI analysis (second and third rows). Because in the initial stages of analysis, all scans from both before and after training are registered to a template derived from their average, the baseline residual Jacobian values in each figure lie close to the mean.

The analysis showed increased coupling between plOFC/AI and left lOFC/12o (Fig. 4, supplementary table 5). Paired t-tests confirmed effects reflected both increases in activity coupling in DisRev learners (t_8_=−3.52, p=0.008) and decreases in controls (t_13_=2.33, p=0.037). For illustration we present the averaged residual timeseries, after controlling for age and gender, extracted from ROIs placed at the center of gravity of the significant cluster (3.375 mm^3^ in diameter) in the DisRev learners and All Controls.

As in the DBM analysis we sought to confirm that the changes in functional coupling were not dependent on non-specific effects but specific to the experience of reversing reward contingencies. We therefore examined the contrast between the DisRev learners and Dis Control learners (after excluding the NoDis Control animals). This more selective analysis confirmed increased coupling between plOFC/AI and left lOFC/12o (Fig. 4C, Supplementary Table 5). For illustration we present the averaged residual timeseries, extracted from bilateral ROIs placed at the center of gravity of the significant clusters in the DisRev learners and Dis Controls.

### Experiment 4

Finally, we examined functional coupling of plOFC/AI in the lesion animals relative to controls (we considered here fMRI data collected at Scan 1). Again a seed-based correlation analysis focused on the same plOFC/AI ROI. The GLM used was equivalent to the one used to examine structural changes in Experiment 2; it sought regions in which functional connectivity was reduced in lesion animals compared to controls. We report reduced coupling between plOFC/AI and lOFC/12o and bilateral ACC/MCC in the lesion group, one cluster in each anatomical ROI (Fig. 4D, Supplementary Table 5). For illustration we present the averaged residual timeseries, extracted from bilateral ROIs placed at the center of gravity of the significant clusters in the 2 lesion animals and the 22 control animals used in this analysis (although structural data were available for 28 control animals, fMRI data were only available for 22 animals).

Table 1 summarizes the DBM results (experiments 1 and 2), resting state fMRI results (experiments 3 and 4) from the learning (experiments 1 and 3) and lesion (experiments 2 and 4) investigations.

**Table 1:**
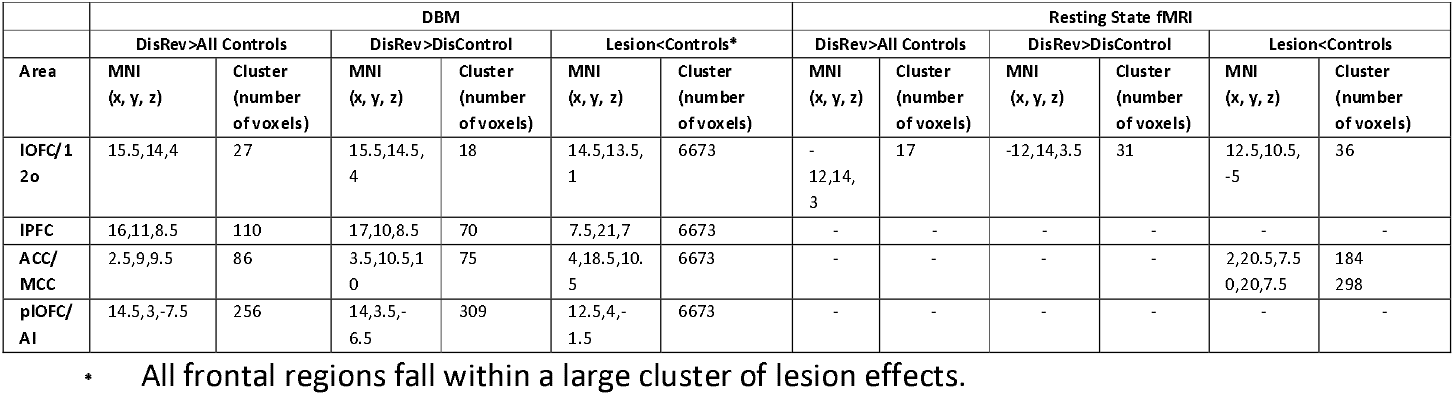
Summary of the results (MNI coordinates and cluster size) across the four analyses (Experiment 1 to 4) and two imaging methods (DBM or resting state fMRI)

## Discussion

Learning to perform the DisRev task proficiently is associated with an extensive and bilateral region of grey matter change in plOFC/AI (experiment 1). Smaller regions of grey matter change were present in three other frontal cortical regions, lOFC/12o, ACC/MCC, and lPFC. However, notably, there were no changes in central OFC. These results complement recent claims that central OFC has only a limited role in DisRev [3, 46]. Moreover, again in-line with the suggestion from Rudebeck and colleagues that aspiration lesions of central OFC, disconnect the regions that are critical for reversal learning, we found that grey matter was reduced in all four of these areas when lesions were made in central OFC (experiment 2). Experiment 3 confirmed that the way in which plOFC/AI interacted with other frontal cortical regions changed when animals learned the DisRev task. Moreover, the patterns of interaction found in experiment 3 were disrupted in experiment 4.

It would be possible to argue for a role of plOFC/AI and the other frontal areas in updating stimulus-outcome associations on the basis of the increased grey matter volume found when DisRev learners were compared to control subjects that did not learn any discrimination task. However, experiments 1 and 3, employed a second analysis in which DisRev learners were compared with control subjects trained to perform very similar two-option reward-guided visual discrimination tasks but which lacked any reversal component. This made it possible to demonstrate that plOFC/AI is especially concerned with reversal learning. Behavioral adaptation in humans has also been associated with activity in AI [47, 48]. The extensive effects found in plOFC/AI are also consistent with a recent finding that activity in the same region carries a signal that reflects not just whether a choice is rewarded but it also reflects the average level of reward received regardless of which choice is taken [49]. It is precisely these two quantities, the history of rewards for specific choices and the average level of rewards regardless of specific choices, which must be considered and balanced in a very particular manner when animals learn DisRev. Unlike in many natural environments, in DisRev, animals must learn to assign more prominence to specific choice-reward associations at the expense of the general reward history.

The co-recruitment of plOFC/AI and another region near to, but outside, central OFC, lOFC/12o, may reflect direct connections between the areas [50, 51]. In rat, pharmacological lesions of the lateral orbital and anterior insula are associated with reversal learning deficits [20]. Future studies will aim at disentangling the specific roles of the two regions in primates, plOFC/AI and lOFC/12o, closest to central OFC in promoting behavioural flexibility. It is also possible to link the present results with fMRI activity recorded in a nearby region when monkeys learn to use specific choice-reward outcomes to guide decision making [9] and reversal learning or task switching in humans [52, 53], reinforcing the idea of a similar structural-functional organization of the OFC and its subdivisions in macaques and humans [54]. Lesions that include this OFC region disrupt the process of reward credit assignment to specific choices [14] while anterior insula reversible lesions are associated with retrieval of goal value to guide decisions [21].

Considerable effort has been dedicated to identifying the specific contributions of OFC to behavior [1–4, 14, 55, 56]. It is becoming increasingly clear that its role can only be understood if allowance is made for functional heterogeneity within it. For example, activity in a reward-guided task varies across different OFC subdivisions [57] and the central OFC appears to have a particularly important role in determining the identity and desirability of rewards, and other aspects of reward-guided decision making assessed by devaluation tasks. By contrast, the present results suggest fluent DisRev performance is mediated by interactions between plOFC/AI and lOFC/12o.

Finally, our results raise the possibility that interactions with other areas within the DisRev network, including ACC/MCC and lPFC are also important. These two regions have been previously associated with reversal learning [14, 58]. ACC/MCC holds information about the value of choices other than the one that is being taken right now and translates such counterfactual choice values into actual behavioral change and exploration [30, 31, 59, 60]. DisRev may also be mediated by the acquisition of cognitive sets or task models, dependent on other interacting brain regions, representing the inter-relationships between the different choice-reward associations active in the task at different times [61, 62]. It is possible that anterior insula is also associated with such processes [63].

In addition to aiding identification of the neural circuit mediating behavioral flexibility, the present results have more general implications. Individual variation in prefrontal circuits has been linked to variation in a wide variety of behavioral measures including general intelligence and lifestyle demographics on the one hand and to psychiatric illnesses on the other hand [64, 65]. The present results, however, suggest inter-individual variability in circuits comprising prefrontal components may not only have an endogenous source. Instead they may also reflect differences in experience that have placed varying demands on cognitive mechanisms in different subjects. Just as the structural changes in our macaques were correlated with improved ability to negotiate the challenges of DisRev so too the neural variation in healthy humans and patients may also be associated with variation in the ability to deal with new cognitive challenges.

## Methods

In total 30 animals (seven females) were involved in the study (Supplementary Table 1).

### Experiment 1: Discrimination reversal learning

Four animals (OB1-4) learned an object-based DisRev task (Object DisRev), five animals (SB1-5) learned a spatial-based DisRev task (Spatial DisRev) and a total of 20 (Control) animals acted as controls (C1-20). See Supplementary information and table S1 for details. Half the twenty control animals (C1-10) had no experience of formal training (NoDis Control), while the other half had experience of performing two-option reward-guided visual discrimination tasks (Dis Control). Crucially, however, no controls had experience of discrimination reversal. Two MRI scans were acquired in all animals, one prior to DisRev training and one after the learning criterion had been met (Supplementary Methods). Data from all of these animals were analyzed in a DBM analysis of brain structural changes associated with discrimination reversal learning. Of these 20 control animals, 14 animals were included as controls in the fMRI analysis. This subset was chosen because they had all received the same isoflurane anesthetic agent as the experimental animals, OB1-4 and SB1-5; the anesthetic agent can impact on fMRI analyses although it does not impact on DBM structural analyses.

### Experiment 2: Central and medial OFC aspiration lesion

Two animals received lesions of central and medial orbitofrontal (OFC) and adjacent ventromedial prefrontal cortex (vmPFC) (LESIONa1, LESIONa2 referred to as vmPFC/OFC lesions; figure 3A) and a comparison of brain structure and functional activity coupling was made between them and a control group (n=28 for structural analysis, n=22 for fMRI analysis). In total 30 macaques were included in the deformation based morphometry (DBM) analysis of brain structural changes and 24 macaques in the resting functional magnetic resonance imaging (fMRI) analysis (again it was not possible to include all animals in the fMRI analysis). Table S1 details demographic information for the aspiration lesion and control animals used in Experiment2. The control animals were OB1-4, SB1-5, C1-10, and C12-20. Both control group and lesion group scans were obtained prior to any learning of discrimination reversal (DisRev) tasks.

### Training Histories

The nine animals in the reversal learning groups (OB1-4, SB1-5) were trained on the following protocol. Animals were initially trained to touch a blue target on screen. The target stimulus appeared either on the left or right of the screen in blocks of 50 trials for a total of 100 trials. The animals had their first scan once they had reached a performance criterion of 90%.

After the first scan the training regime introduced target-based discrimination reversals in an incremental manner. Analogous procedures were used for both the object choice and spatial choice tasks (Object DisRev and Spatial DisRev). The first testing schedule, schedule 1 (Fig. 1) primed the rewarded choice for 25 trials before introducing, in addition, the unrewarded choice on the opposite side of the screen for an additional 75 trials. The choice-reward contingencies were reversed after the animals performed above 85% correct for two consecutive days. The animals experienced two choice-reward reversals under this protocol. The second testing schedule, schedule 2, followed the same pattern but now the rewarded choice was not primed. Animals completed 100 trials where one of two simultaneously presented options was designated as the rewarded target for the day. Again, contingencies reversed after two consecutive days of 85% correct performance. Animals experienced a total of five reversals under this protocol. With perfect performance an animal could, therefore, complete this second phase in ten days. Schedules 3 and 4 introduced a choice-reward association reversal within a given day’s testing session; the reversal occurred once the animal had correctly chosen the target 50 times. The animals completed two days of 100 trials and three days of 150 trials on schedules 3 and 4 respectively. There was no performance criterion in these phases. In the final schedule, the rewarded targets reversed after 25 correctly performed trials. In total the animals had to perform 150 correct trials in a day. Once the animals’ performance was over 80% for two consecutive days, with a subsequent 12 days of consolidation training, their second scan was taken.

The training experience of the twenty control monkeys varied within the group but critically did not include choice reward reversal learning. Four animals had been trained on a fixation task. Six animals were involved in a neuroanatomy study so had no formal training between their two scans [66]. These ten animals are referred to as the No-Discrimination Controls (NoDis Controls). Another ten animals, Discrimination Controls (Dis Controls), learned to discriminate between rewarded objects but had not experienced choice-reward reversals. No surgical intervention occurred between the two scans.

Note that in the first two experiments, two animals (LESION1, LESION2) received lesions of the central and medial OFC and adjacent vmPFC (referred to as a vmPFC/OFC lesion). Prior to their lesion these animals had learned to discriminate between objects but had no experience of choice-reward reversals.

### Apparatus

Animals were trained via positive reinforcement to stay inside a magnetic resonance imaging (MRI) compatible chair in a sphinx position that was placed inside a home-made mock scanner that simulated the MRI scanning environment. They made responses on a touch-sensitive monitor (38 cm wide × 28 cm high) in front of them, on which visual stimuli could be presented (eight-bit color clipart bitmap images, 128 × 128 pixels).

Smoothie rewards (banana mixed with diluted blackcurrant squash) were delivered from a spout immediately in front of the animal. At the end of testing the animals were given their daily food allowance, consisting of proprietary monkey food, fruit, peanuts, and seeds, delivered immediately after testing each day. This food was supplemented by a forage mix of seeds and grains given approximately 6 h before testing in the home cage. Stimulus presentation, experimental contingencies, and reward delivery were controlled by a computer using in-house programs.

### Lesion surgery

At least 12 h before surgery, macaques were treated with an antibiotic [8.75 mg/kg amoxicillin, intramuscularly (i.m.)] and a steroidal anti-inflammatory (20 mg/kg methylprednisolone, i.m.) to reduce the risk of postoperative infection, edema, and inflammation. Additional supplements of steroids were given at 4-to 6-h intervals during surgery. On the morning of surgery, animals were sedated with ketamine (10 mg/kg, i.m.) and xylazine (0.5 mg/kg, i.m.), and given injections of atropine (0.05 mg/kg), an opioid (0.01 mg/kg buprenorphine), and a non-steroidal anti-inflammatory (0.2 mg/kg meloxicam) to reduce secretions and provide analgesia, respectively. The monkeys were also treated with an H2 receptor antagonist (1 mg/kg ranitidine) to protect against gastric ulceration, which might have occurred as a result of administering both steroid and non-steroidal antiinflammatory treatments. Macaques were then moved to the operating theater where they were intubated, switched onto sevoflurane anesthesia, and placed in a head holder. The head was shaved and cleaned using antimicrobial scrub and alcohol.

A midline incision was made, the tissue retracted in anatomical layers, and a bilateral bone flap removed. All lesions were made by aspiration with a fine-gauge sucker. Throughout the surgery, heart rate, respiration rate, blood pressure, expired CO^2^, and body temperature were continuously monitored. At the completion of the lesion, the wound was closed in anatomical layers. Nonsteroidal anti-inflammatory analgesic (0.2 mg/kg meloxicam, orally) and antibiotic (8.75 mg/kg amoxicillin, orally) treatment was administered for at least 5 d postoperatively. All surgery was carried out under sterile conditions with the aid of a binocular microscope. Aspiration lesions of the central and medial orbitofrontal/ventromedial prefrontal cortex were intended to resemble the aspiration lesions that Rudebeck and colleagues[3] found disrupted DisRev task performance; they were therefore placed between the lateral orbitofrontal sulcus and the rostral sulcus (predominantly Walker’s areas 11, 13, and 14).

Approximately four months later the animals were scanned under anesthesia. Using the same protocol as described below (which was the same for control animals) we collected structural and resting state images.

### MRI Data Collection

Protocols for animal care, MRI, and anesthesia were similar to those that we have previously described [35, 36]. During scanning, under veterinary advice, animals were kept under minimum anesthetic levels using Isoflurane. A four-channel phased-array coil was used (Windmiller Kolster Scientific, Fresno, CA). Structural scans were acquired using a T1-weighted MPRAGE sequence (no slice gap, 0.5×0.5×0.5 mm, TR=2,500 ms, TE = 4.01 ms, 128 slices). Whole-brain BOLD fMRI data were collected for 53 min, 26 s from each animal, using the following parameters: 36 axial slices, in-plane resolution 262 mm, slice thickness 2 mm, no slice gap, TR=2,000 ms, TE = 19 ms, 1,600 volumes.

### Deformation Based Morphometric (DBM) Analysis of Structural MRI Data

All the brains were first aligned to the MNI rhesus macaque atlas template [67, 68] using the affine registration tool FLIRT [69, 70] followed by nonlinear registration using FNIRT [71, 72] which uses a b-spline representation of the registration warp field [73]. The resulting images were averaged to create a study-specific template, to which the native grey matter images were then nonlinearly reregistered, creating a 4D image. To avoid potential misalignment concerns, in experiment 1 the lesion animals were not used to create the study-specific template. In experiment 2 all animals (Learners and Controls) were included. We then extracted the determinant of the Jacobian of the warp field used on registered partial volumes to correct for local expansion or contraction from the 4D image. The Jacobian is a matrix of the directional stretches required to register one image to another, and the determinant of this matrix gives a scalar value for the volumetric change implied. The Jacobian values were then used as the dependent variable in the statistical analyses of reversal learning (experiment 1) or the effects of the central OFC and medial OFC/vmPFC aspiration lesion (experiment 2).

General linear model (GLM) analyses were adapted from Winkler and colleagues [34] allowing where necessary permutation inference analysis with our repeated-measure experimental design [two structural scans collected during each of the two scan periods (pre- and post-learning) per subject]. In addition to our regressor of interest (behavioral condition in Experiment 1: All learners verses All Controls or control versus lesion group in Experiment 2) we also included control regressors indexing age and sex of individual monkeys. We implemented the Randomise procedure to perform permutation-based nonparametric testing, examining positive and negative contrasts. The approach was used to identify brain regions which were larger at scan 2 compared with scan 1 in the discrimination reversal learning groups (Object DisRev and Spatial DisRev) compared to Dis Controls only. In experiment 2 we focus on grey matter decrements related to the lesion.

To investigate bilateral grey matter changes we applied an uncorrected threshold of p < 0.032 to both hemispheres and binarised the thresholded image. The right hemisphere image was then dimensionally transformed along the x-axis (around the mid-sagittal plane) and linearly registered with the left hemisphere using FLIRT. The two images were then multiplied allowing us to examine grey matter in areas in which effect significance was p<0.001 and extended over 15 voxels (corresponding to 1.125 mm^3^).

For illustrative purposes Jacobian values were extracted for each structural scan from binarised 3 mm^3^ cube masks placed at the center of gravity of the effect clusters (p=0.001) in both hemispheres. In experiment 1 we present the Jacobian values in the two groups at scan time 1 and 2. In experiment 2 we simply present the Jacobian values in the lesion and control groups separately. The residual variance in Jacobian values, after sex and age were accounted for, was then averaged across hemisphere and structural scans. To represent variance, we index Jacobian values for each contributing animal.

### Functional MRI (fMRI) Analysis of Activity Coupling

Prior to fMRI analysis, the following preprocessing was applied: removal of non-brain voxels, discarding of the first six volumes of each fMRI dataset, 0.1 Hz low-pass filtering to remove respiratory artifacts, motion correction, spatial smoothing (Gaussian 3 mm FWHM kernel), grand-mean intensity normalization of the entire 4D dataset by a single multiplicative factor, high-pass temporal filtering (Gaussian-weighted least-squares straight line fitting, with sigma =50.0 s). Registration of functional images to the skull-stripped structural MRI scan and to the MNI macaque template [67, 68] was achieved with linear and nonlinear registration using FLIRT and FNIRT respectively [69].

To establish changes in functional connectivity within the network as a function of DisRev learning (experiment 1) or lesions (experiment 2) we used a voxel-wise whole brain approach to map resting-state functional connectivity of the plOFC/AI. The 15.6 mm^3^ cube regions of interest (ROI) were made at the center of the grey matter cluster identified in the DBM analyses in experiment 2, displaced minimally to reduce white matter inclusion. The binary images were registered to each monkey’s fMRI scan. The blood oxygen level dependent (BOLD) timeseries was then extracted from each ROI in each individual animal. Analyses were performed using tools from FSL [74] using the method described by Mars and colleagues [75].

For each animal, first, we calculated the first eigen time series of the BOLD signal in the ROI. The first eigen time series is the single time series which best reflects coherent activity across the ROI in that it represents the largest amount of variance across the set of voxels within the region. At the individual subject level, we fitted a GLM consisting of the first eigen time series and seven confound regressors, namely the average time series of the whole brain and six movement parameters expressing movement during the scan as calculated using the FSL tool MCFLIRT (note the scans were obtained from anesthetized animals while their heads were fixed in a stereotaxic frame).

### Experiment 3: Discrimination reversal learning

The resting state analyses were equivalent to those described for the structural-based experiments. A GLM was created with a repeated-measure experimental design [fMRI scan (Scan 1 and Scan 2) per subject]. In addition to our regression of interest for this analysis (All learners (n=9) verses All Controls (n=14)) we also included control regressors indexing age and sex of individual monkeys. Randomise was implemented to perform permutation-based nonparametric testing, examining positive contrasts. Resulting voxelwise p maps were small volumes cluster corrected using threshold-free cluster enhancement methods (p < 0.05) using anatomical masks focused on the frontal regions identified across experiment 1 and 2; namely lOFC/12o (left = 1235 mm^3^, right = 1187.75 mm^3^), lPFC (left = 674.25 mm^3^, right = 614 mm^3^) and ACC/MCC (left = 1222.25 mm^3^, right = 100.75 mm^3^). The same approach was used to identify brain regions which were larger at scan 2 compared with scan 1 in the DisRev learning animals compared to Dis Controls only.

### Experiment 4: Central and medial OFC aspiration lesion

Finally, in experiment 4 we examined plOFC/AI functionally connectivity in Lesion animals (n=2) compared to All Controls (n=22) from their post lesion or pre-learning scans respectively and again sought a between-group difference with reduced connectivity in Lesion compared to Control animals.

For illustrative purposes we present the averaged residual functional connectivity values following the same procedure described for the DMN analysis. For all analyses functional connectivity values were extracted from binarised 3 mm^3^ cube masks placed at the center of gravity of the effect clusters in significant hemispheres. The residual variance in functional connectivity values, after sex and age were accounted for, was then averaged across hemispheres. To represent variance we index functional connectivity values for each contributing animal.

## Acknowledgements

We thank K. Murphy, L. Fisher, E. Bartlett, M. Martinez and Biomedical Services staff for their assistance. JS is funded by a Wellcome Trust Henry Dale Fellowship (105651/Z/14/Z). MN is supported by an Academy of Medical Sciences and Wellcome Trust Springboard fund SBF003\1143 KK, AHB, MEW and MFS are funded by the Wellcome Trust Strategic Award 101092/Z/13/Z. KK is also funded by BBSRC grant BB/H016902/. The work of R.B.M. is supported by the Biotechnology and Biological Sciences Research Council (BBSRC) UK [BB/N019814/1] and the Netherlands Organization for Scientific Research NWO [452-13-015]. The Wellcome Centre for Integrative Neuroimaging is supported by core funding from the Wellcome Trust [203139/Z/16/Z].

