## Supplemental Material for "Behavioral flexibility is associated with changes in structure and function distributed across a frontal cortical network in macaques"

**METHODS**

The work with animals reported in this study was conducted under authority of personal and project license issued by the British Home Office in accordance with the UK Animals (Scientific Procedures) Act (1986).

**Subjects**

In total 30 animals (seven females) were involved in the study (Table S1).

**Table 1**

| Animal | Group | Learning History | Age Scan1 | Age Scan2 | Age Scan3 | Sex |
| --- | --- | --- | --- | --- | --- | --- |
| OB1 | Object DisRev |  | 4.36 | 4.95 | 5.14 | M |
| OB2 | Object DisRev |  | 3.93 | 5.03 | 5.16 | M |
| OB3 | Object DisRev |  | 4.05 | 4.97 | 5.10 | M |
| OB4 | Object DisRev |  | 4.54 | 5.23 | 5.34 | M |
| SB1 | Spatial DisRev |  | 4.40 | 5.14 | 5.39 | M |
| SB2 | Spatial DisRev |  | 4.39 | 5.01 | 5.21 | M |
| SB3 | Spatial DisRev |  | 4.58 | 4.87 | 5.02 | M |
| SB4 | Spatial DisRev |  | 4.69 | 5.06 | 5.29 | M |
| SB5 | Spatial DisRev |  | 4.01 | 5.08 | 5.29 | M |
| C1† | NoDis Control | no task learned | 11.21 | 11.86 | 11.94 | M |
| C2† | NoDis Control | no task learned | 10.47 | 10.93 | 11.07 | M |
| C3 | NoDis Control | visual fixation task | 3.04 | 4.33 |  | M |
| C4† | NoDis Control | no task learned | 5.49 | 5.86 | 5.90 | F |
| C5† | NoDis Control | no task learned | 5.31 | 5.68 | 5.72 | F |
| C6 | NoDis Control | visual fixation task | 3.60 | 4.84 |  | M |
| C7 | NoDis Control | visual fixation task | 2.41 | 4.71 |  | M |
| C8† | NoDis Control | visual fixation task | 3.44 | 4.05 | 4.38 | M |
| C9 | NoDis Control | no task learned | 2.08 | 4.38 |  | F |
| C10† | NoDis Control | no task learned | 3.44 | 4.06 | 4.41 | F |
| C11* | Dis Control | target discrimination task | 3.79 | 6.05 |  | F |
| C12 | Dis Control | target discrimination task | 4.79 | 5.06 |  | F |
| C13 | Dis Control | target discrimination task | 6.97 | 7.27 |  | M |
| C14 | Dis Control | target discrimination task | 6.99 | 7.38 |  | M |
| C15 | Dis Control | target discrimination task | 6.90 | 7.30 |  | M |
| C16 | Dis Control | target discrimination task | 6.49 | 6.86 |  | M |
| C17 | Control | target discrimination task | 4.37 | 4.77 |  | M |
| C18 | Dis Control | target discrimination task | 4.39 | 4.74 | 4.99 | M |
| C19 | Dis Control | target discrimination task | 3.71 | 4.11 | 4.44 | M |
| C20 | Dis Control | target discrimination task | 3.50 | 3.74 |  | M |
| LESIONa1* | OFC/vmPFC lesion | target discrimination task | 6.05 | 6.55 |  | F |
| LESIONa2 | OFC/vmPFC lesion | target discrimination task | 6.15 | 6.69 |  | F |

*C11 and LESIONa1 are the same animal. The data identified as C11 is the animal’s pre-operative scan. While the data from this animal could be used for control purposes in the longitudinal experiment 2 it could not be used as control data in the between-subject experimental design employed in the lesion experiments. †Monkeys scanned with sevoflurane and not isoflurane were not used in the fMRI analysis in experiment 1 or 2 but only in the DBM structural MRI analysis. While the type of anesthetic used may impact on fMRI data it does not impact on structural MRI data.

**RESULTS**

**Structural changes associated with discrimination reversal learning**

**Table 2: DBM results table: Learners > All Controls (experiment 1) Scan 2 > Scan 1**

| Region | x | y | x | Cluster extent (num vox) p < 0.001 |
| --- | --- | --- | --- | --- |
| lOFC (12o) | 15.5 | 14 | 4 | 27 |
| lPFC (46v) | 16 | 11 | 8.5 | 110 |
| IPFC (46d) | 13 | 9.5 | 12 | 54 |
| ACC/MCC (24c) | 6 | 12 | 10.5 | 25 |
| ACC/MCC (24a) | 2.5 | 9 | 9.5 | 86 |
| plOFC/AI | 14.5 | 3 | -7.5 | 256 |
| Striatum (caudate) | 6 | 8 | 7 | 26 |
| Striatum (caudate) | 4.5 | 1 | 8.5 | 30 |
| Striatum (putamen) | 9.5 | 4.5 | 0.5 | 42 |
| Basal Forebrain (Basal Nucleus of Meynert) | 4 | 2.5 | -4.5 | 116 |
| Inferotemporal cortex (TE) | 16.5 | 0.5 | -14.5 | 26 |
| Inferotemporal cortex (TEO) | 27 | -13.5 | 3 | 45 |
| Posterior STS (Tpt) | 21.5 | -16 | 9 | 21 |
| Amygdala (ABmc) | 7.5 | 0 | -7.5 | 87 |
| Hippocampus | 7.5 | -4.5 | -9 | 48 |
| Substantia Nigra | 4.5 | -10 | -8 | 28 |
| Parietal cortex (7b) | 20 | -13.5 | 13.5 | 75 |
| Occipital Cortex (V4) | 14 | -19.5 | -3 | 29 |
|  | 18.5 | -24.5 | -5.5 | 26 |
| Occipital Cortex (V4) | 22.5 | -25 | 8 | 83 |
|  | 26 | -24.5 | 3.5 | 45 |
|  | 25 | -24.5 | -4.5 | 71 |
| Occipital Cortex (V3) | 19.5 | -28.5 | -6.5 | 146 |
|  | 9.5 | -31 | 17 | 1995 |
| Occipital Cortex (V2) | 26 | -24 | 0.5 | 15 |
|  | 8.5 | -29.5 | 12 | 43 |
|  | 5.5 | -31.5 | 4.5 | 33 |
|  | 7 | -28.5 | 3 | 35 |
| Occipital Cortex (V1) | 25.5 | -28 | -0.5 | 82 |
|  | 25 | -29.5 | 1.5 | 39 |
|  | 23 | -32.5 | -0.5 | 18 |
| Cerebellum | 10.5 | -26.5 | -1.5 | 23 |
|  | 2.5 | -24 | -0.5 | 85 |
|  | 7.5 | -32 | -1 | 49 |

**Table 3: DBM results table: Learners > DisControls (experiment 1) Scan 2 > Scan 1**

| Region | x | y | z | Cluster extent (num vox) p < 0.001 |
| --- | --- | --- | --- | --- |
| mOFC (14r) | 2 | 18.5 | -1.5 | 20 |
| lOFC (12o) | 15.5 | 14.5 | 4 | 18 |
| lPFC (46v) | 18.5 | 14.5 | 7.5 | 15 |
| lPFC (8Ad/v) | 17 | 10 | 8.5 | 70 |
| ACC/MCC (24c) | 6 | 16 | 12 | 20 |
| ACC/MCC (24c) | 5.5 | 15 | 9.5 | 45 |
| ACC/MCC (24b) | 3.5 | 10.5 | 10 | 75 |
| plOFC/AI | 14 | 3.5 | -6.5 | 309 |
| Premotor Cortex (F2) | 7 | 9.5 | 18 | 16 |
| Striatum (caudate) | 5.5 | 7.5 | 6.5 | 55 |
| Striatum (caudate) | 5 | 2 | 8.5 | 83 |
| Basal Forebrain (Nucleus Basalis of Meynert) | 3.5 | 1.5 | -5 | 31 |
| Striatum (putamen) | 10 | 3 | -0.5 | 136 |
| Insula (Id) | 19 | 0.5 | -1.5 | 21 |
| Inferotemporal Cortex (TGa) | 16.5 | 1.5 | -15 | 102 |
| Entorhinal cortex | 9 | -3 | -14.5 | 18 |
| Posterior STS | 23.5 | -12 | 4.5 | 54 |
|  | 18 | -13.5 | -0.5 | 25 |
|  | 26.5 | -14 | 3 | 18 |
| Amy (ABmc) | 6 | -1.5 | -8 | 52 |
| Hippocampus | 7.5 | -4 | -8.5 | 31 |
| PCC (23b) | 3.5 | -12 | 12.5 | 40 |
| Parietal cortex (7b) | 20 | -13.5 | 13.5 | 23 |
| Parietal cortex (PEa) | 11 | -19.5 | 21.5 | 52 |
| Parietal cortex (PEc) | 5 | -25.5 | 22 | 207 |
| IPS (LIP) | 10.5 | -25.5 | 21.5 | 16 |
| Occipital Cortex (V2/V3) | 5 | -32.5 | 15.5 | 177 |
|  | 3.5 | -28 | 18.5 | 17 |
| Occipital Cortex (V4) | 14 | -27.5 | 19.5 | 146 |
| Occipital Cortex (V3) | 11 | -26.5 | 12 | 53 |
|  | 10.5 | -28.5 | 15.5 | 24 |
| Occipital Cortex (V2/V3) | 9.5 | -32.5 | 16 | 64 |
| Occipital Cortex (V2) | 23.5 | -32 | -0.5 | 28 |
|  | 9.5 | -37 | 11 | 115 |
|  | 8.5 | -33 | 19.5 | 71 |
| Cerebellum | 1.5 | -23.5 | 0 | 16 |
|  | 19 | -28.5 | -7 | 50 |

**Structural changes associated with lesions**

**Table 4: DBM results table: Lesion study (experiment 2)**

| Region | x | y | z | Cluster extent (num vox) p < 0.001 |
| --- | --- | --- | --- | --- |
| lPFC (12r/46v) | 10.5 | 24.5 | 6.5 | 17 |
| lOFC (12o) | 14.5 | 15 | -2 | 16 |
| MPFC (8Bm) | 2 | 14 | 17.5 | 42 |
| cOFC (11/13) | 7 | 15 | 2 | 6673 |
| Somatosensory cortex (SII) | 22.5 | -2 | 2 | 43 |
| Inferotemporal (TEa) | 14 | 3.5 | -15 | 84 |
| Inferotemporal cortex (TEm) | 26 | -12.5 | -1.5 | 32 |
| mid-STS (PGa) | 17.5 | -1 | -8 | 22 |
| STG | 24.5 | -5 | -5 | 99 |
| Posterior STS | 12.5 | -24.5 | 17.5 | 76 |
| Amygdala (Lv) | 10.5 | -1 | -12.5 | 42 |
| Hippocampus | 15 | -10.5 | -6.5 | 49 |
| PCC (23) | 4 | -13.5 | 9 | 42 |
| Parietal cortex (7a) | 17 | -26 | 16.5 | 21 |
|  | 21 | -18.5 | 14 | 27 |
| Intraparietal sulcus (VIP) | 8.5 | -18.5 | 12 | 74 |

**Functional changes associated with OFC lesions and discrimination reversal learning (experiment 3 and 4)**

**Table 5: fMRI-measured activity coupling bilateral Results table:**

| Contrast | Region | x | y | z | Cluster extent (num vox) p < 0.001 |
| --- | --- | --- | --- | --- | --- |
| Dis Rev Learners > All Controls (experiment 2) Scan 2 > Scan 1 | lOFC (12o) | -12 | 14 | 3 | 17 |
| DisRev learners > Dis Controls (experiment 2) Scan 2 > Scan 1 | lOFC (12o) | -12 | 14 | 3.5 | 31 |
| OFC Lesion animals < All Controls | lOFC (12o) | 12.5 | 10.5 | -5 | 36 |
|  | Rostral ACC (right) | 2 | 20.5 | 7.5 | 184 |
|  | Rostral ACC (left) | 0 | 20 | 7.5 | 298 |
|  | ACC/MCC (left, 24) | -1.5 | 12.5 | 9.5 | 20 |
